# Codon usage is a stochastic process across genetic codes of the kingdoms of life

**DOI:** 10.1101/066381

**Authors:** Bohdan B. Khomtchouk, Claes Wahlestedt, Wolfgang Nonner

**Affiliations:** Department of Psychiatry and Behavioral Sciences, University of Miami Miller School of Medicine, 1120 NW 14th Street Suite 1463, Miami, FL 33136, USA; Department of Physiology and Biophysics, University of Miami Miller School of Medicine, 1600 NW 10th Avenue, Miami, FL 33136, USA

## Abstract

DNA encodes protein primary structure using 64 different codons to specify 20 different amino acids and a stop signal. To uncover rules of codon use, ranked codon frequencies have previously been analyzed in terms of empirical or statistical relations for a small number of genomes. These descriptions fail on most genomes reported in the Codon Usage Tabulated from GenBank (CUTG) database. Here we model codon usage as a random variable. This stochastic model provides accurate, one-parameter characterizations of 2210 nuclear and mitochondrial genomes represented with > 10^4^ codons/genome in CUTG. We show that ranked codon frequencies are well characterized by a truncated normal (Gaussian) distribution. Most genomes use codons in a near-uniform manner. Lopsided usages are also widely distributed across genomes but less frequent. Our model provides a universal framework for investigating determinants of codon use.

## Introduction

Determining a mathematical law followed by codon rank distributions of biological species would be useful for investigating the nature of the evolutionary processes that have acted upon these distributions throughout time (1). A universal, i.e., biological species independent, distribution law for codons has remained elusive for almost three decades (1, 6–14).

The ‘language’ by which genomes describe proteins has received theoretical interest ever since the genetic code was discovered. In particular, a degeneracy of vocabulary is intriguing: the 64 different codons of the DNA genetic code outnumber the different amino acids and stop signal that they encode by a factor ≈ 3. How do biological organisms deal with or exploit such degeneracy? Does biased use of synonymous codons encode information beyond amino acid sequence (2–6)? Information flowing from genome to encoded proteins can be monitored at its source by counting how often certain codons are used by a genome. Normalizing codon counts into frequencies and ordering frequencies into a descending series generate frequency-rank plots that are comparable across genomes in spite of differences in genome size, genetic code, or varying bias in the use of synonymous codons. Such plots provide a global perspective for investigative and theoretical work regarding the organization of the coding DNA and the machineries of translation.

Toward a mathematical interpretation of frequency-rank plots, various empirical formal descriptions have been proposed: a power of rank (Zipf’s law) (7, 8), an exponential of rank (9–11), or a combination of exponential and linear relations (1). Also, various statistical relations have been formulated: the first statistical model of codon use was made by Borodovsky et al (12, 13) and, more recently, by Naumis et al (14, 15). Most of these descriptions do not capture the tail of codon/rank plots that have an inflection, a feature observed in many genomes. Models based on additive or multiplicative contributions to codon use have been devised to better describe the tail of such frequency-rank plots (1, 14, 15).

In this paper we present a systematic study of codon use based on the Codon Usage Tabulated from GenBank (CUTG) database (16). Codon frequency is interpreted as a random variable, and the ranks of codon frequencies are interpreted in terms of cumulative probability. Cumulative probability is described by a truncated normal distribution, which is used in our analysis as an empirical description rather than as the consequence of a specific stochastic model. We analyze codon use in 2210 genomes throughout the CUTG database. Differences in codon use among diverse genomes are captured in a single adjustable parameter. We establish a near-continuous variety of codon use that gravitates strongly toward uniform use of codons in all kingdoms of life.

## Methods

CUTG database entries comprising at least 10^4^ codons are analyzed. Observed relative codon frequencies *y* of an organism are ordered so that rank *r* = 1 is given to the largest frequency: *y*(*r*), 1 ≤ *r* ≤ 64. The discrete rank-frequency series *r*(*y*) is interpreted in terms of the continuous rank-frequency function

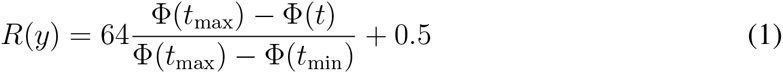

where Φ is the normal distribution in the standardized variable *t* = (*y* − *μ*)/*σ*, that is, the cumulative probability of events in the range −∞ through *t*. That distribution is naturally truncated here to the range of possible codon frequencies, 0 ≤ *y* ≤ 1, or *t*_min_ = −*μ*/*σ* through *t*_max_ = (1 − *μ*)/*σ*. We normalize the truncated distribution, complement, and map linearly to ranks to construct the continuous rank function *R*(*y*), which we will superimpose to the discrete rank series of observed codon frequencies, *r*(*y*). We rotate the conventional frequency/rank plot (e.g., Fig 1B, C) counterclockwise by 90 degrees, so that frequency becomes the independent variable (abscissa) and rank the dependent variable (ordinate, proportional to the cumulative probability of codon frequency).

**Fig 1.**
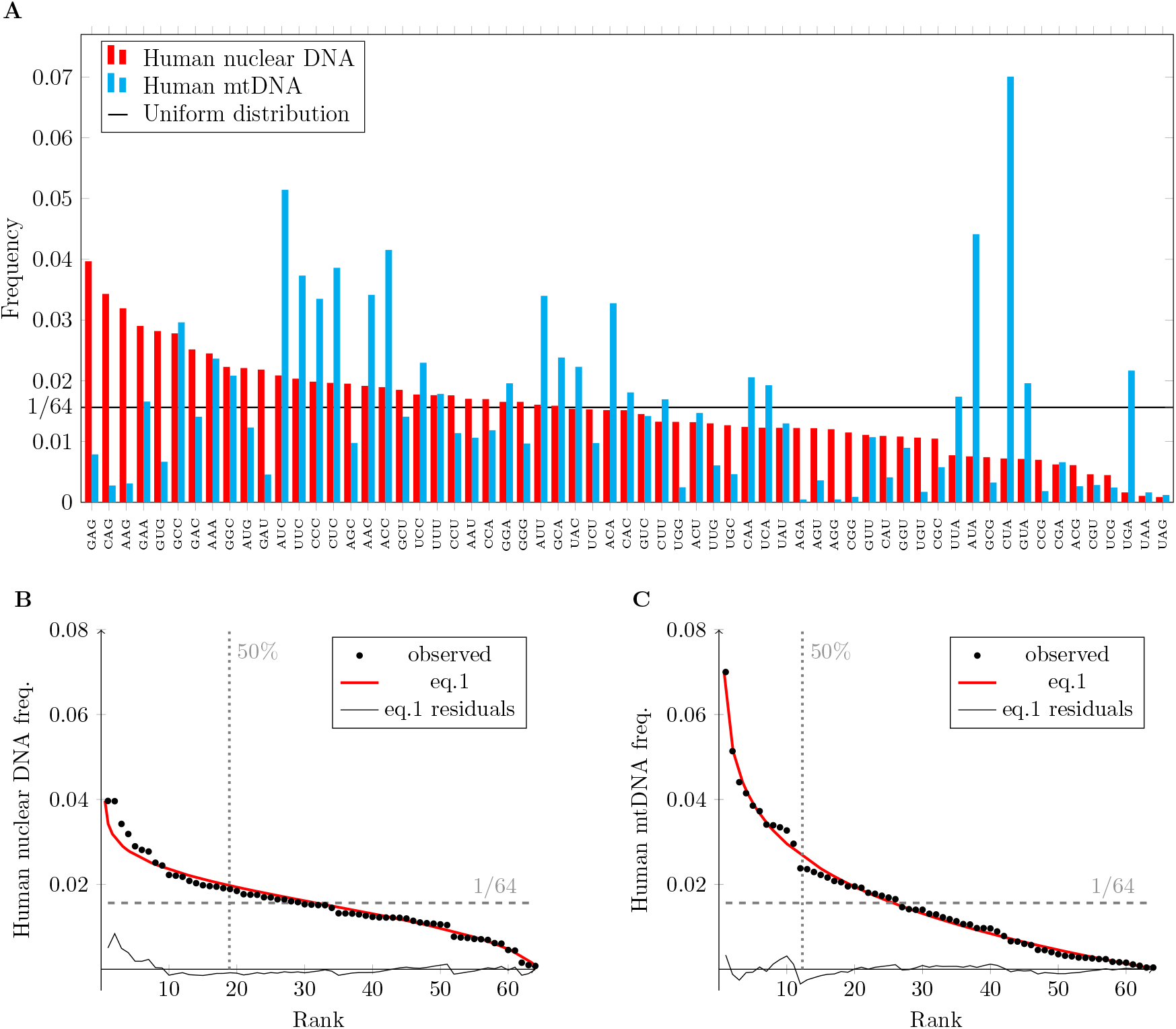
Human codon use. (A) Relative codon frequencies of the nuclear and mitochondrial DNA. Codons are ranked to give a descending order for the nuclear DNA. (B) and (C) Stochastic model superimposed to ranked frequencies of nuclear and mitochondrial DNA, respectively. (For model parameters see file S1 in Supplementary Materials.)

Since the observed frequencies *y_i_* are normalized (∑_*i*_ *y_i_* = 1), the average codon frequency is 1/*N*_codons_ = 1/64. A truncated distribution to be superimposed to these observed frequencies thus must have a mean of 1/64. In the case of the truncated normal distribution

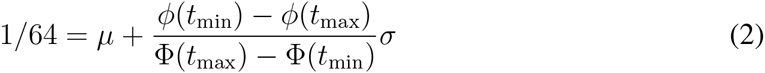

where *φ*(*t*) = *d*Φ(*t*)/*dt*. Note that location μ and scale *σ* also enter the relation through the definition of *t*.

Since frequency normalization establishes a relation between the normal-distribution parameters *μ* and *σ*, these cannot be chosen independently in fitting *R*(*y*) (eqn. 1) to the observed ranks *r*(*y_i_*). Only one of these parameters is free. We vary *σ* to minimize ∑_*i*_(*r*(*y*_i_) − *R*(*y_i_*))^2^ while computing the value of *μ* to be adjoined to a choice of *σ* from eqn. 2.

## Results

In interpreting ranked frequencies of codon occurrence one must keep in mind that ranking abstracts the distribution of frequencies from the identities of the codons. Codons occupying particular frequency ranks in one genome typically occupy different ranks in other genomes. Consider the human nuclear and mitochondrial genomes (Fig. 1A). Both sets of codon frequencies are ranked here in the order of decaying occurrence in the *nuclear* genome. The mitochondrial genome evidently does not follow the nuclear ranking. The mitochondrial codons need to be arranged in a much different order to form a decaying sequence. If the codon frequencies of these genomes are ranked, the same rank generally corresponds to different codons of the genetic code.

When the nuclear and mitochondrial codon frequencies of Fig. 1A are individually ranked in decaying order they reveal two different patterns of codon use (symbols in Fig 1B, C). The nuclear human codons are used with quite uniform frequencies, as evident at middle-ranks, and fewer are used substantially more often or rarely at low and high ranks, respectively (Fig. 1B). The 19 lowest-ranked codons receive about one half of the total usage. In the mitochondrial genome, the 12 lowest-ranked codons receive about one half of the total usage (Fig. 1C). Mitochondrial codon usage is less balanced among codons than nuclear codon usage.

The red lines in Fig. 1B, C represent frequency-rank relations computed from the model that we present in this paper. Codon frequency is interpreted here as a random variable (rather than a probability), and the ranks associated with these frequencies as (unnormalized) estimates of the cumulative probabilities of the frequencies. We fit that cumulative distribution by a truncated Gaussian distribution (eqn. 1). The model reproduces the two different forms of human codon use. These are determined by a single free parameter, the location *μ* of the Gaussian distribution. The other parameter of that distribution, the scale *σ*, is fixed by the parameter *μ* and the requirement that the predicted frequency distribution be normalized (eqn. 2).

Fig. 2 illustrates the differences between the two truncated normal distributions underlying the curves in Fig. 1B, C. Fig. 2A, C show as red lines the cumulative normal distributions, Φ(*t*)/(Φ(*t*_max_) − Φ(*t*_min_)), and Fig. 2B, D the underlying probability density functions, *φ*(*t*)/(Φ(*t*_max_) − Φ(*t*_min_)). The solid black lines are derived from the observed nuclear (Fig. 2A, B) or mitochondrial codon frequencies (Fig. 2C, D) and represent (*r*(*y*) — 0.5)/64 (Fig. 2A, C) or number of codons per bin width of the binned codon usage *y*, respectively (Fig. 2B, D). The range of the truncated distribution for the nuclear codon frequencies comprises the position *μ* of the normal distribution, whereas the range of the mitochondrial frequency distribution is restricted to the right-tail of the normal distribution.

**Fig 2.**
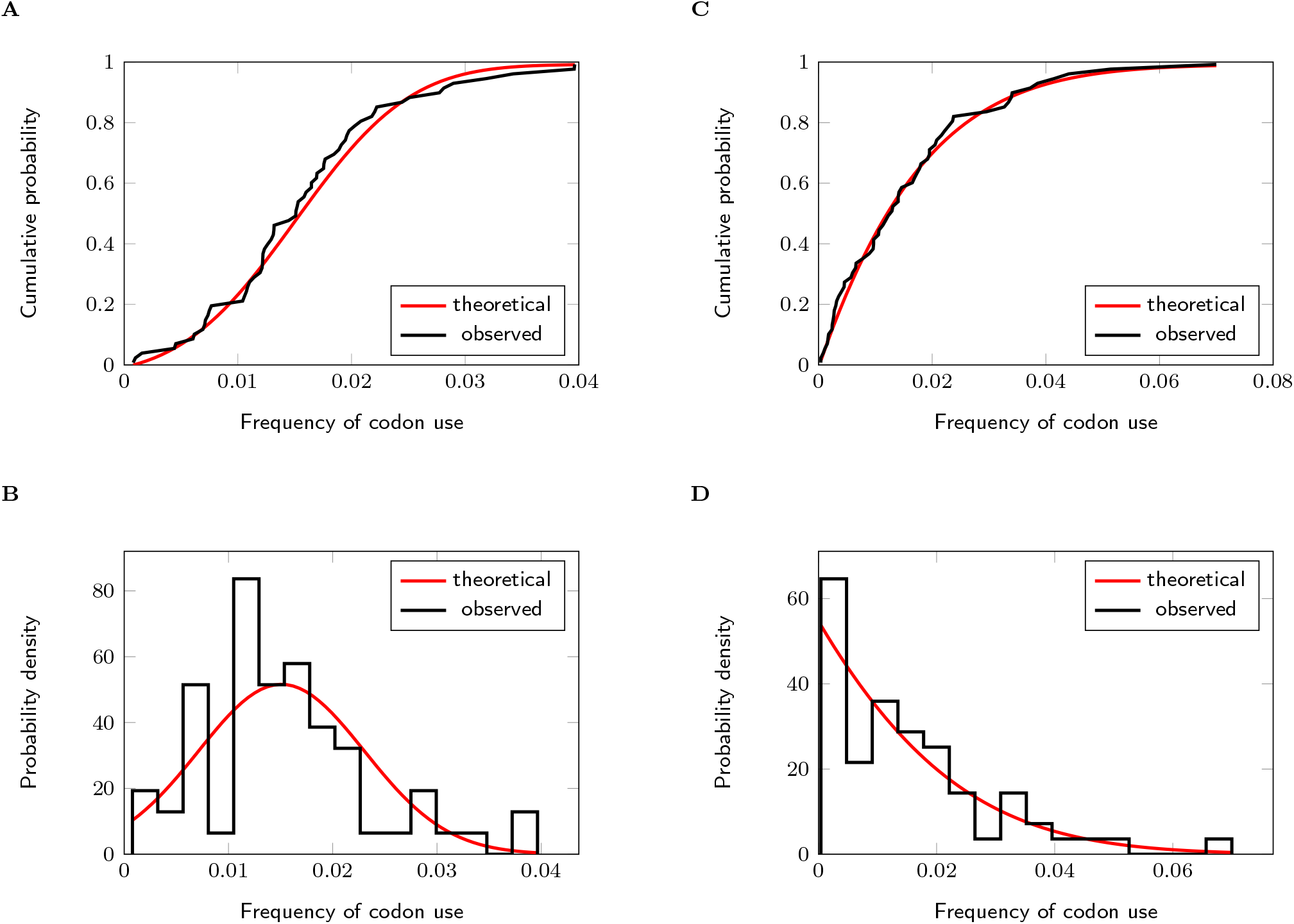
Normal distributions of codon frequency in human nuclear (A, B) and mitochondrial DNA (C, D). For details see text.

We apply the truncated normal distribution model to several genomes that are common subjects of study (Fig. 3, symbols). The model reproduces the varying patterns of codon usage in these genomes (red lines). The parameters for all theoretical curves in Figs. 1 and 3 are compiled in file S1 of Supplementary Materials. Residuals between observed frequencies and the truncated Gaussian model tend to be largest in the most frequently used codons if codon usage is relatively uniform (Fig. 1B; see also Fig. S2 C, D, F–I). A slightly skewed (rather than normal) distribution function might better describe these cases.

**Fig 3.**
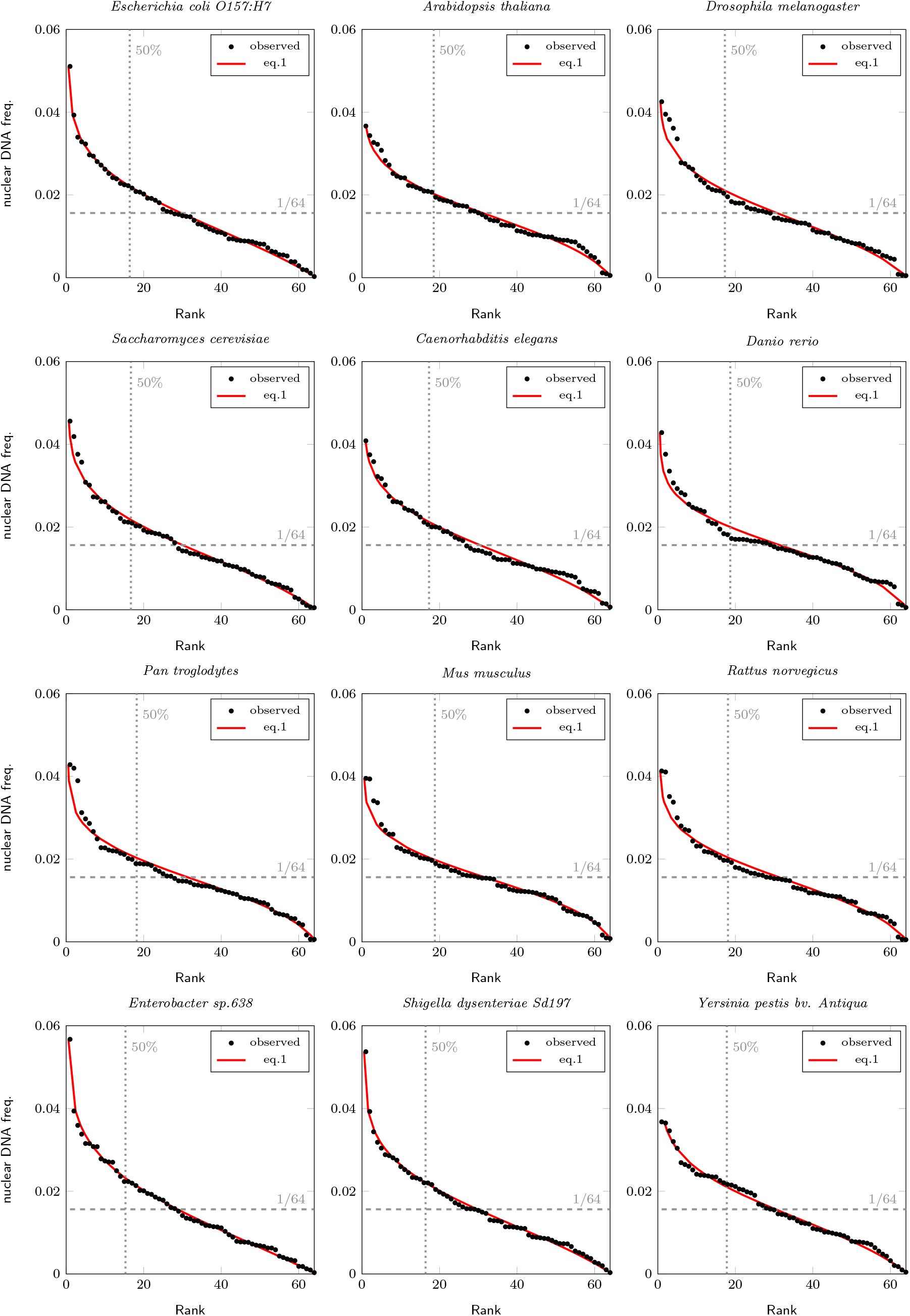
Codon use in nuclear DNA of several species fitted to the stochastic model. (For model parameters see file S1 in Supplementary Materials.)

Since the genomes sampled in Figs. 1 and 3 suggest that codon usage varies substantially among genomes, we analyze genomes throughout the CUTG database with the only restriction being that an entry comprise at least 10^4^ codons. The parameters of the truncated normal distributions describing the nuclear genome entries are summarized in Fig. 4A, and those of mitochondrial genome entries in Fig. 4B. These graphs show both location *μ* and scale *σ* to illustrate the variation of both parameters (*μ* and σ are connected through eqn. 2). A histogram of the parameters *μ* (Fig. 4C) shows that the most preferred pattern of codon use is the more uniform usage (the location of the normal distribution then approaches the mean of the truncated distribution, 1/64, because the peak of the normal distribution is within the truncated range). Both nuclear and (the fewer available) mitochondrial genomes of at least 10^4^ codons, however, also visit a range of lopsided codon usages: the values of *μ* describing these genomes extend far into the negative range so that codon frequency distributes over the right-tail of the normal distribution. Over this full range of observed codon usages, a single parameter, *μ*, characterizes codon frequency distributions and thereby provides a universal and concise means of characterization.

**Fig 4.**
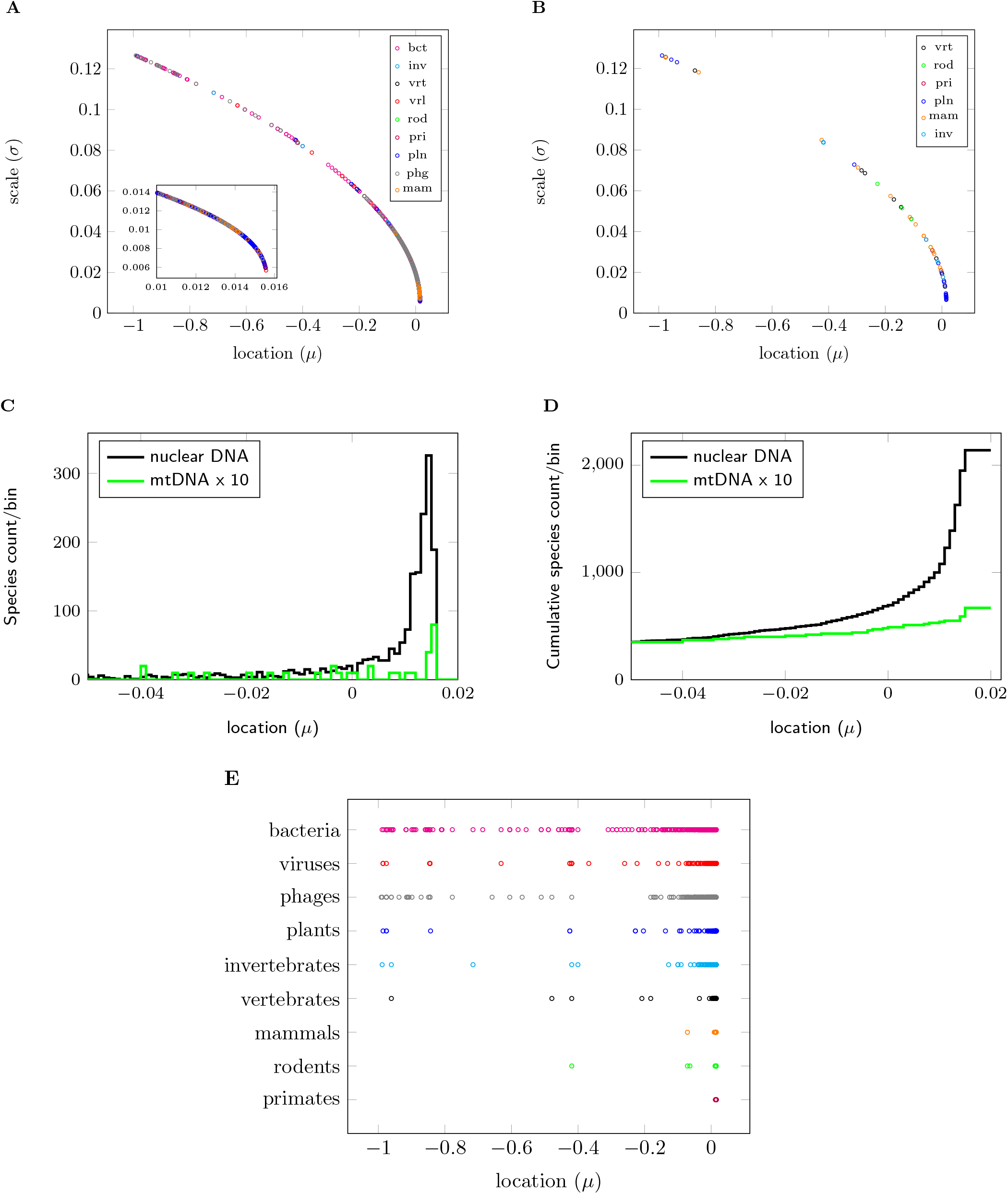
Analysis of codon use across the genomes listed in the CUTG database. Parameters of the truncated normal distributions describing the ranked codon frequencies of each nuclear (A) and mitochondrial genome (B). Histogram (C) and cumulative histogram (D) of the Gaussian parameter *μ*. Values of Gaussian parameters *μ* in each division of the CUTG database (E).

To assess variations of codon usage among and within different kingdoms of life we summarize the values of the parameter *μ* found for nuclear genomes by database division (Fig. 4E). A joint cumulative histogram of *μ* is shown in Fig. 4D. Positive values of *μ* prevail in all divisions – relatively uniform distributions of codon frequency are the rule. In bacteria, phages and viruses asymmetrical codon use (indicated by negative values of *μ*) is also widely distributed. Mitochondrial genomes (Fig. 4B) reveal a similar pattern. The multicellular organisms of the plant and invertebrate divisions also show many occurrences of lopsided codon use, whereas the species in the vertebrate, mammal, rodent, and primate divisions use codons uniformly with few exceptions. A full report for all database entries included in our survey is available in a text file for each division (Supplementary Materials, files S2-S10).

## Discussion and Perspectives

We show here that the ranked frequencies of codon occurrence of a large number of genomes are well characterized by truncated Gaussian distributions with a single adjustable parameter. With regard to the accuracy of description and number of required parameters, we improve substantially on several previously proposed approaches (Fig. S1 and S2).

A statistical relation proposed by Naumis et al (13, 14) also describes codon usage better than other previously proposed models. Despite an appealing quality of fit, however, the Naumis et al two-parameter model does not reveal an underlying systematic structure within the codon usage data. A scatterplot of the empirically determined values of the model parameters *a* and *b* shows an unstructured relationship as if the Naumis et al model fits the data in an ad-hoc manner (Fig. S3A and S3B). Our model gives a comparable fit with a single free parameter (Fig. S2). Fig. S3C and S3D show how the Naumis et al model predicts the different human nuclear and mitochondrial codon frequencies by variations of two factors. Each of these factors controls aspects of predicted codon use over the full range of ranks and is independently controlled by a model parameter, *a* or *b*. A model structured this way might produce similar frequency/rank relations with diverse combinations of values for *a* and *b*. The apparently random variation of these parameters as seen in Fig. S3A, B thus might be due to small inter-genomic differences that produce strong variations among these parameters. Indeed, variations of codon use between lopsided and uniform that are evident in the genomic data do not map to a pattern in the Naumis et al model parameters.

The majority of genomes that we have analyzed are well characterized by a normal distribution of codon frequency that is positioned close to average codon use, 1/64. Such a pattern of codon usage ressembles a snapshot of 64 random variables that follow approximately uniform statistics. Individual codon frequencies in a genome summarize the encoding of many evolving polypeptides and of evolving choices of codons within a degenerate genetic code. Moreover, each CUTG database entry for a species comprises genomes of several specimens. The analyzed codon frequencies thus acquire a stochastic nature. A minority of genomes exhibit codon frequencies that we empirically describe by normal distributions positioned away from average codon use, 1/64. In these cases, the statistics of the individual codon frequencies must be nonuniform. Within the range of codon usage patterns between uniform and lopsided we observe a strong preference toward uniformity. Factors promoting uniform codon usage thus are a natural target for more detailed investigation. Such factors might be universal as they appear to be at work throughout the kingdoms of life.

## Conclusions

We have conducted a cross-species, cross-kingdom analysis of biological organisms in 2210 nuclear and mitochondrial genomes to elaborate universal features in the relative usage of codons. We have found that codon frequency distributions are well described by a truncated normal distribution. Observed patterns of codon usage varying between uniform and lopsided are characterized by a single adjustable parameter. Nuclear genomes universally gravitate toward uniform use of codons. Our stochastic model, and the universal properties of codon usage it reveals, should prove useful in investigating the nature of the evolutionary processes that have shaped codon usage.

## Figure Legends

**Fig S1.**
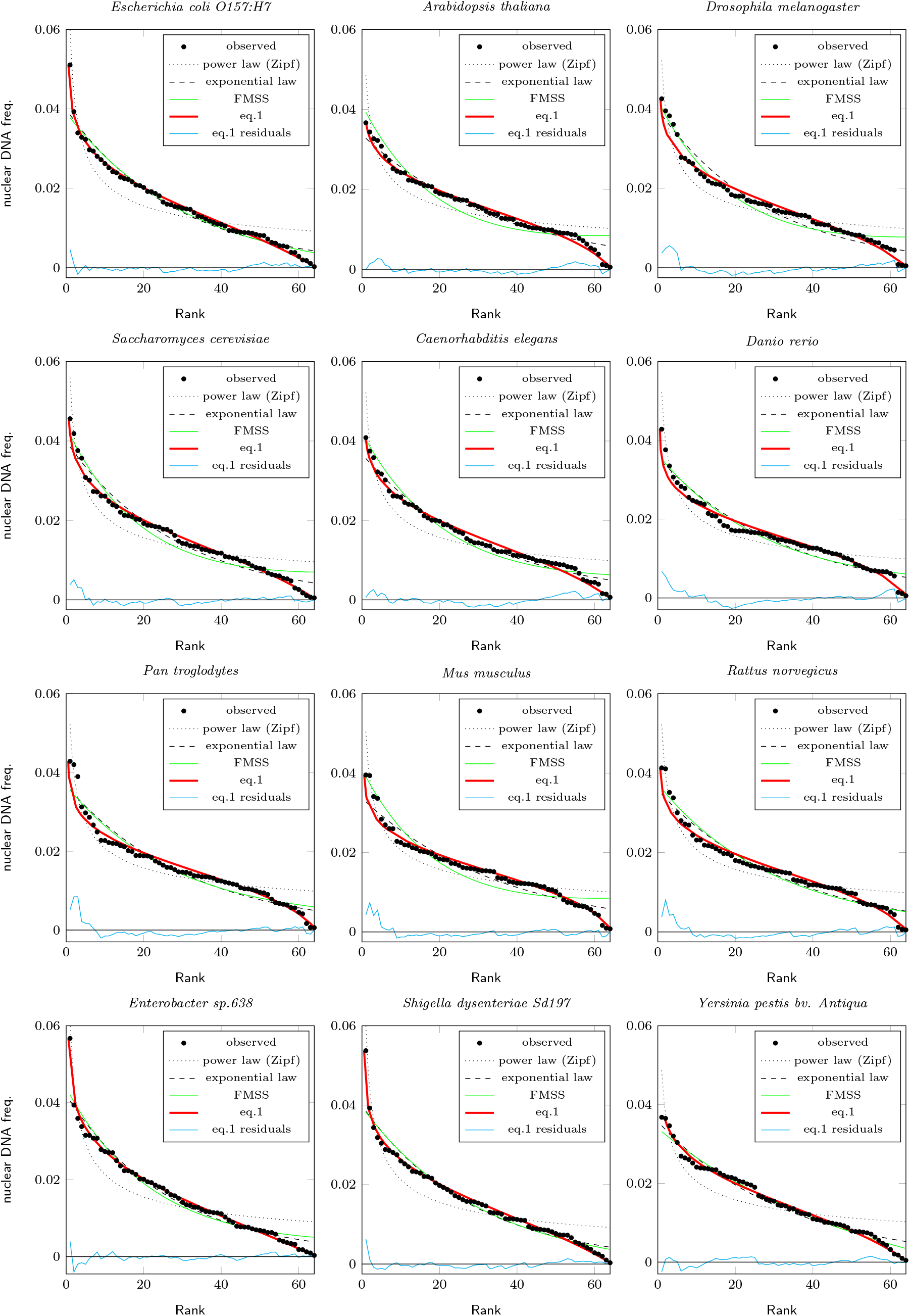
Empirical relations vs. the stochastic model superimposed to ranked frequencies of nuclear DNA. Power law (Zipf): *y*(*r*) = *αr*^−*β*^; exponential law: *y*(*r*) = *α* exp(−*βr*); FMSS (1): *y*(*r*) = *α* exp(−*ηr*) − *βr* + *γ*; Stochastic: eqn. 1. (For model parameters see file S1 in Supplementary Materials.)

**Fig S2.**
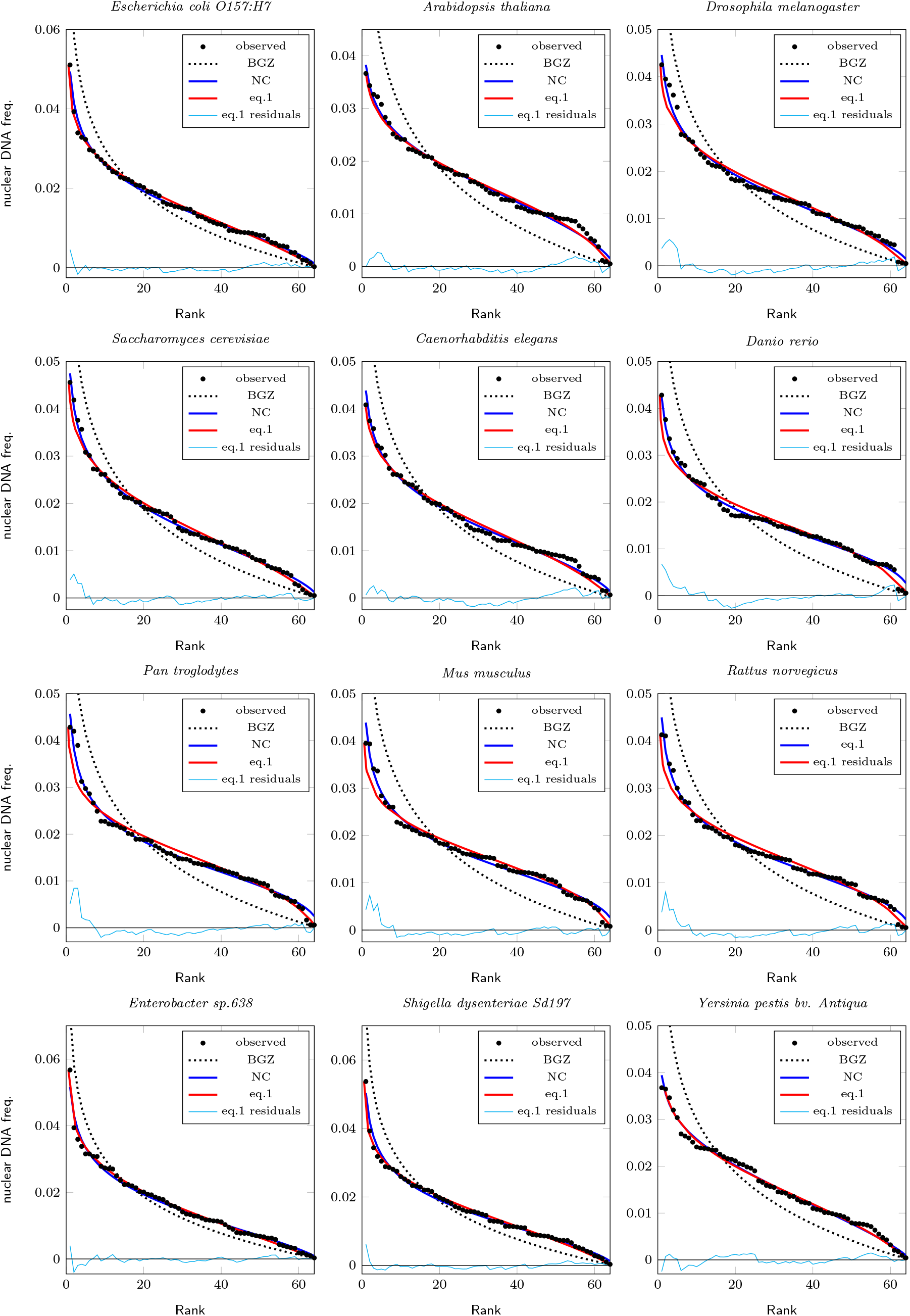
Statistical relations vs. the stochastic model superimposed to ranked frequencies of nuclear DNA. BGZ (11): 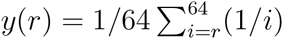; NC (13): 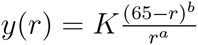; Stochastic: eqn. 1. (For model parameters see file S1 in Supplementary Materials.)

**Fig S3.**
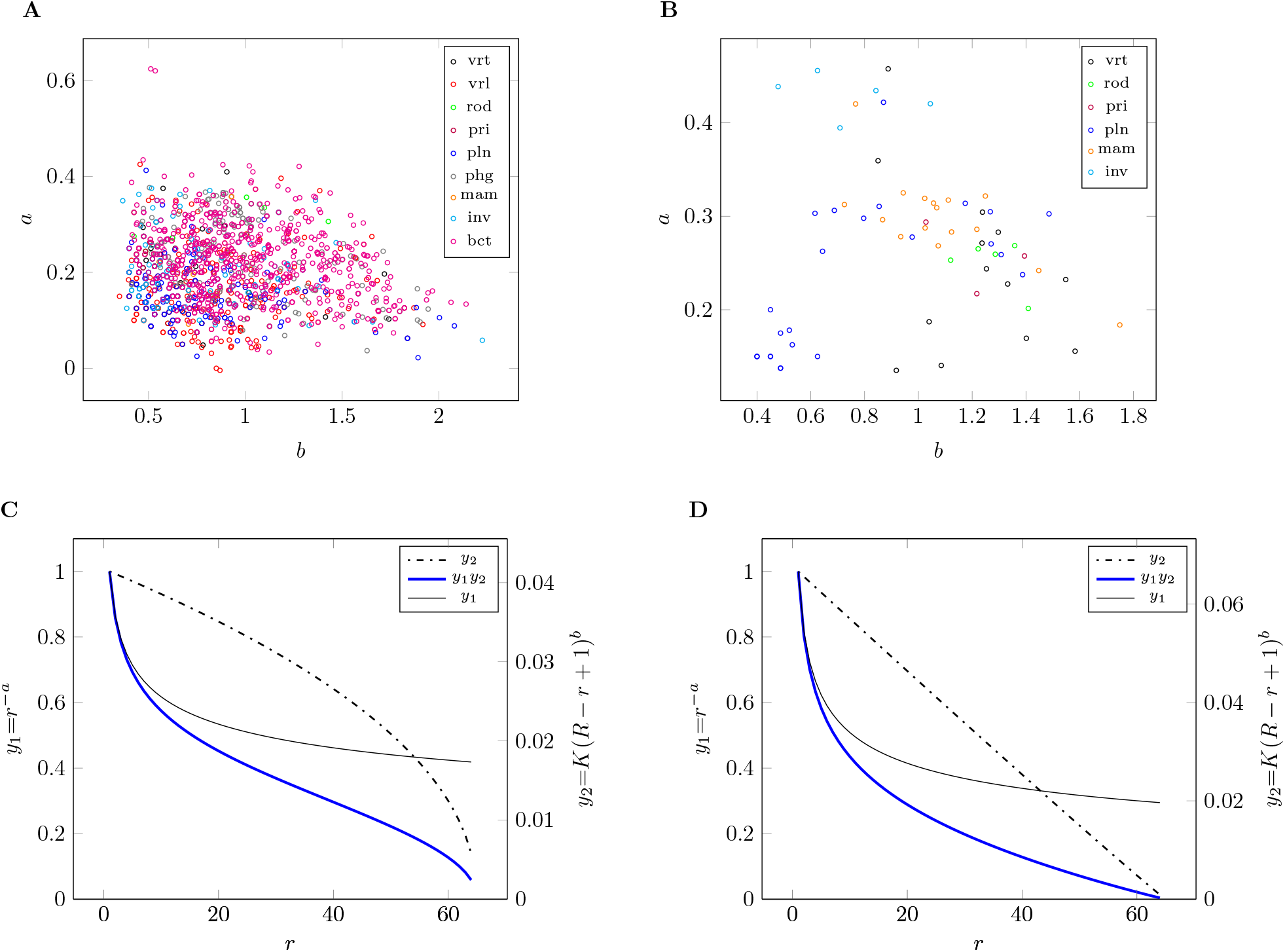
Scatterplot of NC model parameters for nuclear (A) and mitochondrial genomes (B) across kingdoms. Numerator, denominator, and composite NC model plotted for human nuclear (C) and mitochondrial (D) genome.

## Acknowledgments

BBK wishes to acknowledge full financial support by the United States Department of Defense (DoD) through the National Defense Science and Engineering Graduate Fellowship (NDSEG) Program: this research was conducted with Government support under and awarded by DoD, Army Research Office (ARO) Biosciences Division, National Defense Science and Engineering Graduate (NDSEG) Fellowship, 32 CFR 168a.

